# Characterisation of cell cycle checkpoint kinases in Toxoplasma gondii

**DOI:** 10.1101/2024.09.10.612042

**Authors:** Monique K. Johnson, Sara Chelaghma, William H. Lewis, Ludek Koreny, Ross F. Waller, Catherine J. Merrick

## Abstract

*Toxoplasma gondii* is a protozoan parasite in the apicomplexan phylum. Apicomplexan parasites replicate using a variety of non-canonical cell division modes, distinct from binary fission, whose molecular regulation is incompletely understood. *T. gondii* replicates by endodyogeny in its intermediate hosts, and by endopolygeny in its definitive host. To improve our understanding of how these unusual, flexible cell division modes are regulated, we characterised the *T. gondii* homologues of the cell-cycle checkpoint kinases ATM and ATR. These phosphoinositol-3-kinase-like kinases are entirely absent in some related parasites including *Plasmodium*; in *T. gondii* they are present but their putative checkpoint roles were previously uncharacterised. Both *Tg*ATM and *Tg*ATR were found to be dispersed throughout the parasite, rather than restricted to the nucleus, and they did not detectably relocate to the nucleus after DNA damage. Nevertheless, they were both required for checkpoint responses to DNA damage, including acute replication slowing and phosphorylation of the DNA damage marker histone H2AX. Unusually, the two kinases seemed to cooperate in the checkpoint response, with the loss of either one largely ablating the response, regardless of the type of DNA damage. Thus, *T. gondii* clearly retains a DNA-damage-responsive checkpoint, but some of its key features differ from the well-studied checkpoint in human cells.

**AUTHOR SUMMARY:** *Toxoplasma* is one of the most widespread parasites of humans, with a large proportion of the global population being infected. Infection usually causes no symptoms, but can have severe consequences in pregnant women and immunocompromised people. The *Toxoplasma* parasite is a single-celled organism, related to other parasites that cause important human diseases like malaria. All these parasites are distinctive in having unusual modes of replication: unlike human cells, they do not replicate by simply splitting in two. Furthermore, drugs to treat toxoplasmosis usually act to suppress the parasite’s replication. Therefore, it is important to understand how this unusual replication is controlled, both for our fundamental understanding of cell division modes, and to ensure effective drug treatment of toxoplasmosis. Here, we characterised for the first time the proteins that control the *Toxoplasma* cell cycle by responding to DNA damage. We report that a damage-responsive checkpoint exists, but has some unusual features.

## INTRODUCTION

The apicomplexan parasite *Toxoplasma gondii* is a significant cause of disease in both animals and humans. It is widespread in the global human population as a largely asymptomatic infection, but it can cause lethal disease in immunocompromised patients, retinal scarring and blindness in immunocompetent individuals, and severe birth defects if pregnant women are acutely infected. Accordingly, effective drugs are required to treat acute toxoplasmosis, and concerns have recently been raised about resistance to current therapies [1]. These therapies usually consist of anti-folate regimes such as pyrimethamine/sulfadiazine [2], which act to suppress folate metabolism, starve the parasite of nucleotide precursors and suppress its replication. A detailed understanding of the parasite’s mode of replication is therefore important – both to understand the action of anti-parasite drugs, and also because *Toxoplasma* replication is fundamentally unusual. As such, it can illuminate our understanding of the basic biology of eukaryotic cell division.

*Toxoplasma* lies in the apicomplexan phylum of protozoan parasites and a key feature of this deeply-diverging phylum is that the species therein do not replicate by conventional binary fission – the cell division mode used by most model cells from *S. cerevisiae* to human. Instead, they employ a variety of syncytial division modes, such as endodyogeny in *Toxoplasma*, endopolygeny in *Sarcocystis* and schizogony in the malaria parasite *Plasmodium* [3, 4]. Many of the regulatory proteins that drive and control the *Toxoplasma* cell cycle are now being identified [5–9], and some elements of this cell cycle are clearly unusual, such as the apparent separation between the nuclear division and budding cycles [10, 11], but comparatively little is known about stress-responsive checkpoints within this cycle.

Eukaryotic cells make highly conserved ‘checkpoint’ responses to DNA damage or stalled replication, involving a kinase cascade that arrests the cell cycle and promotes DNA repair [12]. This is initiated by one or more members of the phosphoinositol-3-kinase-like kinase (PIKK) family. These kinases – Mitosis entry checkpoint 1 (Mec1) in budding yeast; Ataxia-Telangiectasia Mutated or Rad3-Related (ATM and ATR) in humans – have a catalytic domain that is structurally related to PI3K lipid kinases [13]. However, they act upon protein substrates instead, including the checkpoint-transducing kinases Rad53/Chk1/Chk2 and the histone H2AX, which marks damaged DNA for repair.

As an obligate intracellular parasite, *Toxoplasma* can certainly suffer DNA damage through inflammatory stress, oxidative and free-radical damage, etc. Like most eukaryotes, it encodes pathways to repair DNA double-strand breaks (DSBs): homologous recombination and non-homologous end-joining [14]. Therefore, it would make evolutionary sense to retain DNA-damage-responsive checkpoints. Curiously, these may be absent in at least some apicomplexans: *Plasmodium* species seem to entirely lack PIKK kinases [15], and may not arrest their cycle after aberrant genome replication [16, 17]. *Toxoplasma*, by contrast, does encode PIKK homologues, although the downstream kinases Chk1/2 have not been clearly identified and their DNA damage checkpoint has not been thoroughly investigated, despite its potential importance as a target for anti-parasite drugs [18].

Here, we characterised for the first time the two PIKKs in *T. gondii*, which are annotated *Tg*ATM and *Tg*ATR, according to their homology with human ATM and ATR. Both genes were categorised as ‘fitness conferring’ in a Crispr-based gene knockout screen [19]. To our knowledge, *Tg*ATR was entirely unstudied, while *Tg*ATM was first identified in 2010 [20] in a study of the acetyltransferase *Tg*MYST. This found that *Tg*MYST may induce a putative ATM homologue, *Tg*ATM, and also inferred that this homologue may phosphorylate *Tg*H2AX, although *Tg*ATM was not knocked down or isolated to show this directly. A slow-growth phenotype when *Tg*MYST was over-expressed was taken to represent *Tg*ATM-induced checkpoint activity, and this was ablated by an inhibitor of human ATM, KU-55933, which was assumed (but not directly shown) to inhibit *Tg*ATM as well. A subsequent study also used this inhibitor and showed that KU-55933 itself could slow *T. gondii* growth, and also abolish H2AX phosphorylation after DNA damage [21].

Hence, there was circumstantial evidence that *Tg*ATM acts as DNA damage checkpoint kinase, and could be essential for replication by endodyogeny, but neither *Tg*ATM nor *Tg*ATR had been genetically manipulated or studied directly. Here, we report that *Tg*ATM and *Tg*ATR can both be knocked down without any fitness defect in cultured parasites, but that both proteins – in a largely overlapping manner – do enforce a DNA damage checkpoint in *T. gondii*.

## RESULTS

### *T. gondii* encodes homologues of cell cycle checkpoint kinases

To verify the presence of ATM and ATR orthologues in *T. gondii*, we performed a phylogenetic analysis of known ATM/ATR proteins and their closest matches in *T. gondii* and throughout apicomplexans and other major eukaryotic lineages (Figure 1, S1). This analysis also included the related TOR proteins and the PI3K/PI4K kinases, which formed an outgroup. We identified single *T. gondii* homologues of ATM, ATR and TOR, each of which was most closely related to the respective human homologue. The *T. gondii* homologues of ATM, ATR and TOR were all annotated in ToxoDB as putative forms of these three proteins; the single homologues of the lipid kinases PI3K and PI4K were annotated in ToxoDB only as ‘PI3/4 kinases’ but one gene clearly grouped with PI3Ks, and the other gene with PI4Ks, from both human and *Plasmodium* species.

**Figure 1:**
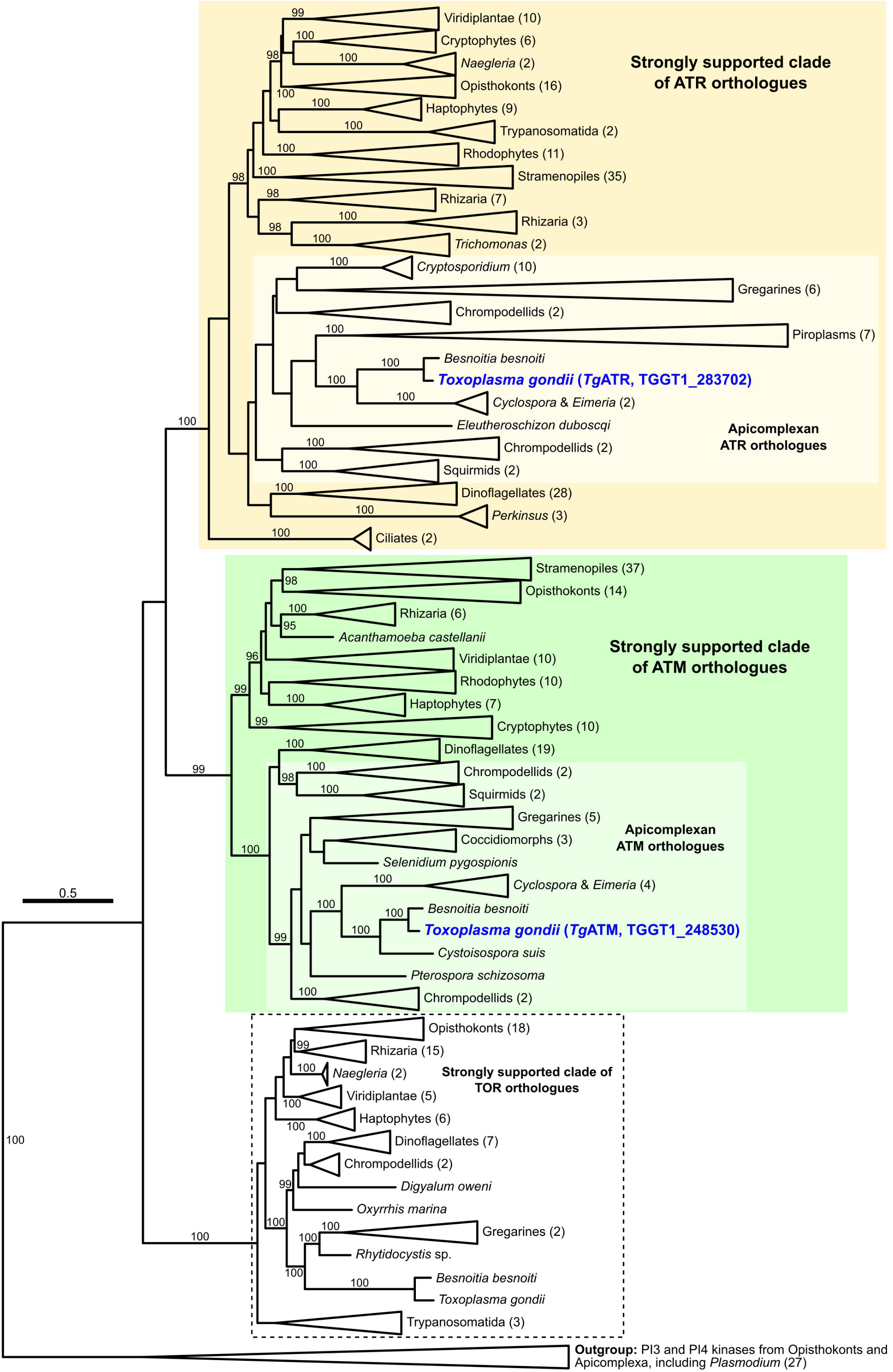
*T. gondii* encodes homologues of cell cycle checkpoint kinases. Phylogeny inferred from amino acid sequences of ATR, ATM and TOR homologues from various eukaryotes. The tree is rooted on PI3K and PI4K sequences, which are paralogues of ATR, ATM and TOR, from Opisthokonts and Apicomplexa. Support values, displayed as percentages, were generated from 1000 ultrafast bootstrap replicates [50] and only support values ≥ 95, indicating that strongly supported clades are shown. Strongly supported clades are highlighted for ATR (support value = 100) and ATM (support value = 99) sequences (yellow and green, respectively). The scale bar represents the number of amino acid substitutions per site. A fully-expanded version of this tree with no collapsed clades and displaying sequence accessions is provided in Figure S1.

The *T. gondii* proteins each grouped, with strong support, with those of other apicomplexans and further related myzozoans, indicating that each of these molecules were present in the common ancestor of all apicomplexans. Conspicuously, *Plasmodium* spp. lacked orthologues for ATM, ATR or TOR, with the nearest orthologues found as outgroups to these PIKKs, i.e. PI3K and PI4K (Figure 1). Similarly, there was independent loss of ATM from *Cryptosporidium* and the piroplasms *Babesia* and *Theileria*. Thus, while both ATM and ATR occur in *T. gondii*, there is an apparent trend towards loss or shift of function of these molecules in Apicomplexa.

In the PIKK family, ATM in human cells responds primarily to DNA DSBs, while ATR responds primarily to stalled replication forks [22]. TOR has a different cell-cycle role, regulating cell growth and protein production. Therefore, in *T. gondii*, homology would suggest that *Tg*ATM and *Tg*ATR should be most directly involved in DNA-damage checkpoint responses to DNA DSBs and stalled replication forks respectively. However, neither protein had yet been investigated, localised in the parasite cell, characterised for kinase activity, or knocked out to determine essentiality.

### Inducible knockdown of *Tg*ATM and *Tg*ATR

PIKKs are very large proteins: *Tg*ATM has a predicted size of ∼250kDa, and *Tg*ATR, ∼650kDa. Cloning for overexpression or recombinant expression would be prohibitively difficult, as would biochemical purification of these proteins. We therefore addressed the function of *Tg*ATM and *Tg*ATR by making inducible knockdowns, since a genome-wide screen for essential genes in *T. gondii* had previously suggested that both genes may be refractory to complete knockout [19] (Figure S2A, B).

The strategy for gene tagging and inducible knockdown is shown in Figure 2A and B: 5’ integration of a haemagglutinin (3xHA) tag and a tetracycline-repressible promoter [23, 24]. PCR genotyping of the resultant mutants confirmed correct integration (Figure S2C, D). Upon induction of knockdown with anhydrotetracycline (ATc), western blotting showed that all detectable HA-tagged *Tg*ATM was lost within 24h and this depletion was stable through 72h (Figure S2E). A similar assessment of the *Tg*ATR knockdown was not possible because the protein was too large for effective western blotting. However, immunofluorescence assays (IFA) showed the near-total loss of both proteins within 24h (Figure 2C, D) and this was confirmed by quantifying the immunofluorescent signals. Residual fluorescence after knockdown was similar to the levels seen in parental cells without any HA-tagged protein (Figure 2E, F).

**Figure 2:**
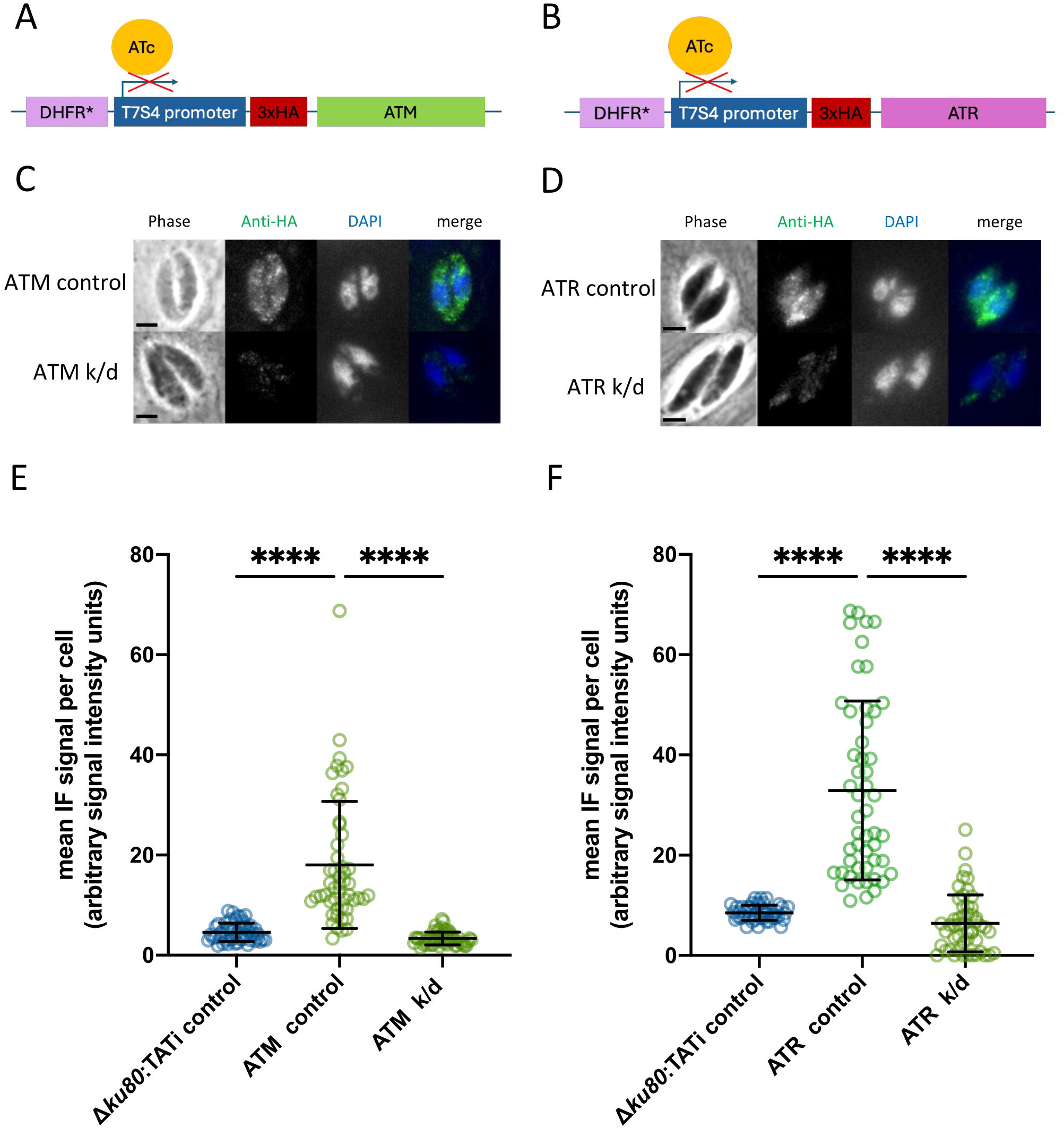
Inducible knockdown of *Tg*ATM and *Tg*ATR. A,B. Constructs used for tagging the *Tg*ATM (A) and *Tg*ATR (B) genes in the *T. gondii* genome. DHFR*, dihydrofolate reductase (selectable marker gene). C,D. IFA showing HA-tagged *Tg*ATM (C) and *Tg*ATR (D) in intracellular tachyzoites: representative cells are shown from each tagged line, before and after addition of ATc for 24 h. Scale bar, 2 μm. E,F Quantification of IFA signals in the parental control line (Δ*ku80*:TATi), each tagged line, and each knockdown (k/d, ATc treated for 24 h): *Tg*ATM (E) and *Tg*ATR (F). N = 50 parasites quantified per condition. Ordinary one-way ANOVA and post hoc Tukey’s multiple comparisons tests of all samples; P values: *<0.05, **<0.01, ***<0.001, ****<0.0001. Comparisons not shown, ns.

Human ATM and ATR have a primarily nuclear location, consistent with their function [25]. Interestingly, however, the reporter-tagged forms of both *Tg*ATM and *Tg*ATR were detected throughout the cell (Figure 2C, D), their diffuse appearance was inconsistent with localisation to any subcellular organelle and confocal microscopy did not reveal further details beyond this generally diffuse location. It is notable that *Tg*MYST, which has been proposed to acetylate *Tg*ATM, was also reported to be cytoplasmic, contrary to its expected nuclear location [20]. Neither *Tg*ATM nor *Tg*ATR appeared in the published subcellular atlas of the *Toxoplasma* proteome [23], and the difficulty of handling such large, unstable proteins precluded subcellular fractionation for western blotting to conclusively determine the proportions of *Tg*ATM and *Tg*ATR in nuclear and cytoplasmic fractions.

### Knockdown of *Tg*ATM or *Tg*ATR imposes no growth defect or DNA-damage-sensitivity in culture

Contrary to predictions from the genome-wide screen [19], neither gene knockdown showed an overt growth phenotype in triplicate plaque assays (Figure 3A, B). We therefore concluded that *Tg*ATM and *Tg*ATR were either entirely dispensable for parasite growth in normal culture conditions, or that these proteins were essential but that residual protein after knockdown, albeit undetectable via western blot or immunofluorescence, could suffice under normal conditions. In either case, challenging the parasites with DNA damage might reveal an effect on parasite growth. Therefore, parasites were subjected to two DNA damaging agents, the alkylating agent methyl methane sulphonate (MMS) or the topoisomerase inhibitor camptothecin (CPT) (previously validated to cause H2AX phosphorylation in this system [21]).

**Figure 3:**
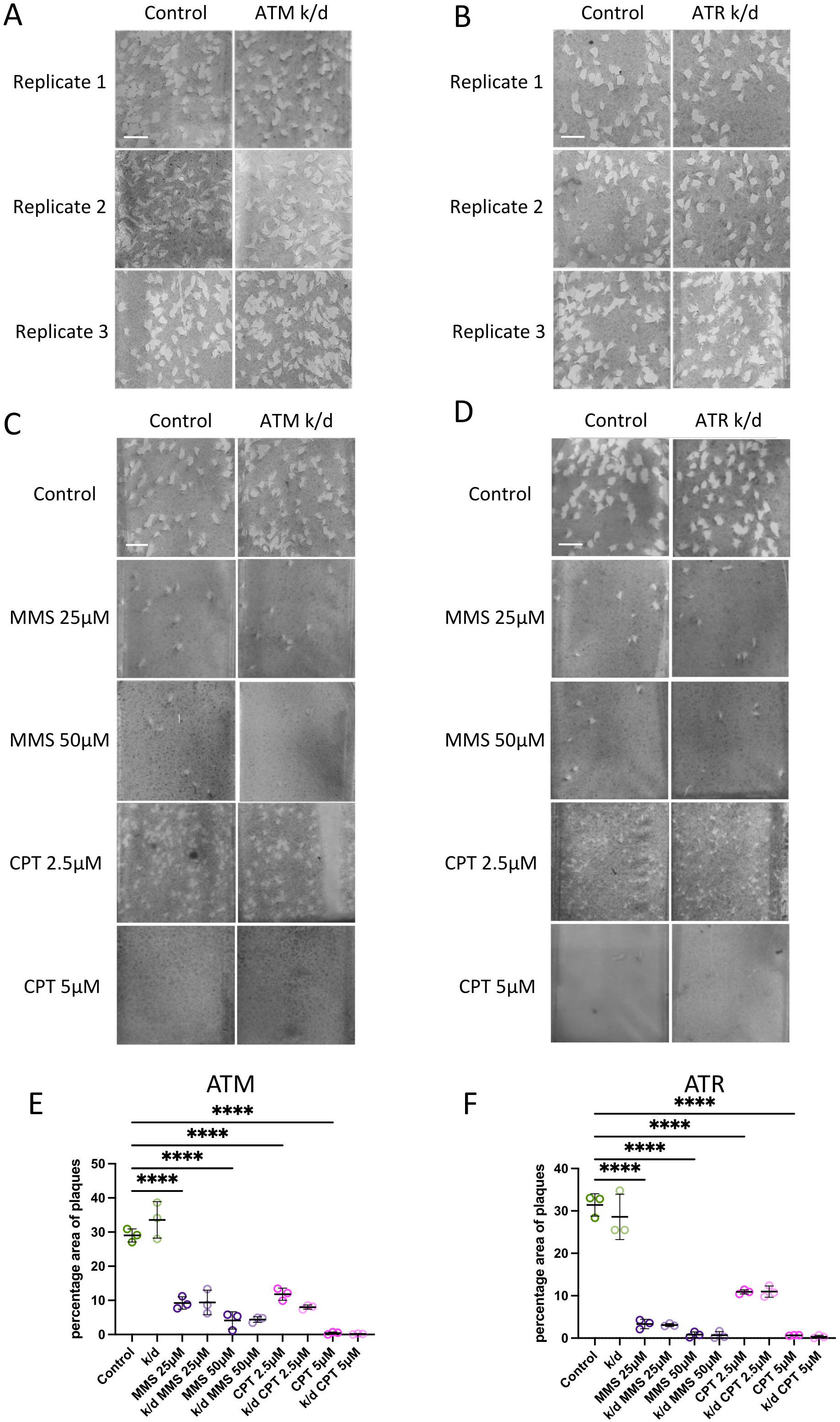
Knockdown of *Tg*ATM or *Tg*ATR imposes no growth defect or sensitivity to DNA damage. A,B. Images of three replicate plaque assays, either untreated (control) or treated with ATc to induce *Tg*ATM k/d (A) and *Tg*ATR k/d (B). C,D. Plaque assay images are shown with or without DNA damage, CPT or MMS, at the concentrations stated for 10 days. *Tg*ATM k/d (C) or *Tg*ATR k/d (D) parasites were pre-treated with ATc for 24 h and then additionally treated with DNA damaging agents for 10 days. Representative images from three biological replicates are shown. E,F. The percentage area of plaques in each background of host cells was quantified from representative images shown in A, B, C and D: three biological replicates were quantified for each condition. Ordinary one-way ANOVA and post hoc Tukey’s multiple comparisons tests were made of each condition vs PIKK k/d, and control vs all; P values: *<0.05, **<0.01, ***<0.001, ****<0.0001. Comparisons not shown, ns.

Both agents caused a titratable defect in parasite growth, the nature of which appeared to be different for MMS versus CPT. MMS primarily reduced the number of plaques, suggesting acute parasite killing followed by a rapid decay of its alkylating activity, so that surviving parasites grew quite normally over the subsequent 10 days. CPT reduced the number but also, primarily, the size of plaques that grew after 10 days, suggesting ongoing growth-inhibitory activity as well. However, none of these effects was clearly exacerbated by the knockdown of either PIKK (Figure 3C-F).

### Knockdown of *Tg*ATM or *Tg*ATR ablates acute checkpoint-like responses to DNA damage

We then reasoned that long-term growth assays might not be a suitable read-out if the knockdowns did, in fact, impede a DNA damage checkpoint. Parasites that failed to respond to DNA damage and continued to replicate, rather than inducing DNA repair, might initially have a growth advantage but might eventually die if their genome was irreparably damaged. Hence the outcome of a 10-day plaque assay might appear similar whether a checkpoint was induced or not. Accordingly, we sought more acute phenotypes in the knockdown parasites.

A 24h ‘replication assay’ quantifies parasite growth during just the first ∼3 cycles of endodyogeny after invasion, by counting the numbers of parasites in newly formed vacuoles in individual host cells (Figure 4A). Notably, this does not establish whether the cells have all arrested in a particular stage – pre-replication (‘G1’) or during S-phase – but simply tests for an acute slowing of their overall cycling rate. This assay confirmed that replication rates were indeed normal in both knockdown lines. DNA damage induced by either MMS or CPT could reduce the initial replication rate and the knockdown of either PIKK ablated this (Figure 4B, C), implying that cell-cycle slowing after DNA damage was indeed the result of checkpoint induction, and that knockdown parasites failed to induce such a checkpoint. There was little difference between the *Tg*ATM and *Tg*ATR knockdowns, suggesting substantial cooperation between the two proteins, with both being similarly required for an effective checkpoint response to two types of DNA damage: alkylation or topoisomerase inhibition. Of note, however,the *Tg*ATM knockdown only partially restored replication to normal rates after CPT damage, but did fully restore it after MMS damage at 25μM, and almost fully after 50μM, while the *Tg*ATR knockdown only partially restored replication to normal rates after MMS damage, but did fully restore it after CPT damage. This suggests that there is some degree of specialisation between the two proteins according to different types of DNA damage, but both proteins were clearly required for a fully effective response.

**Figure 4:**
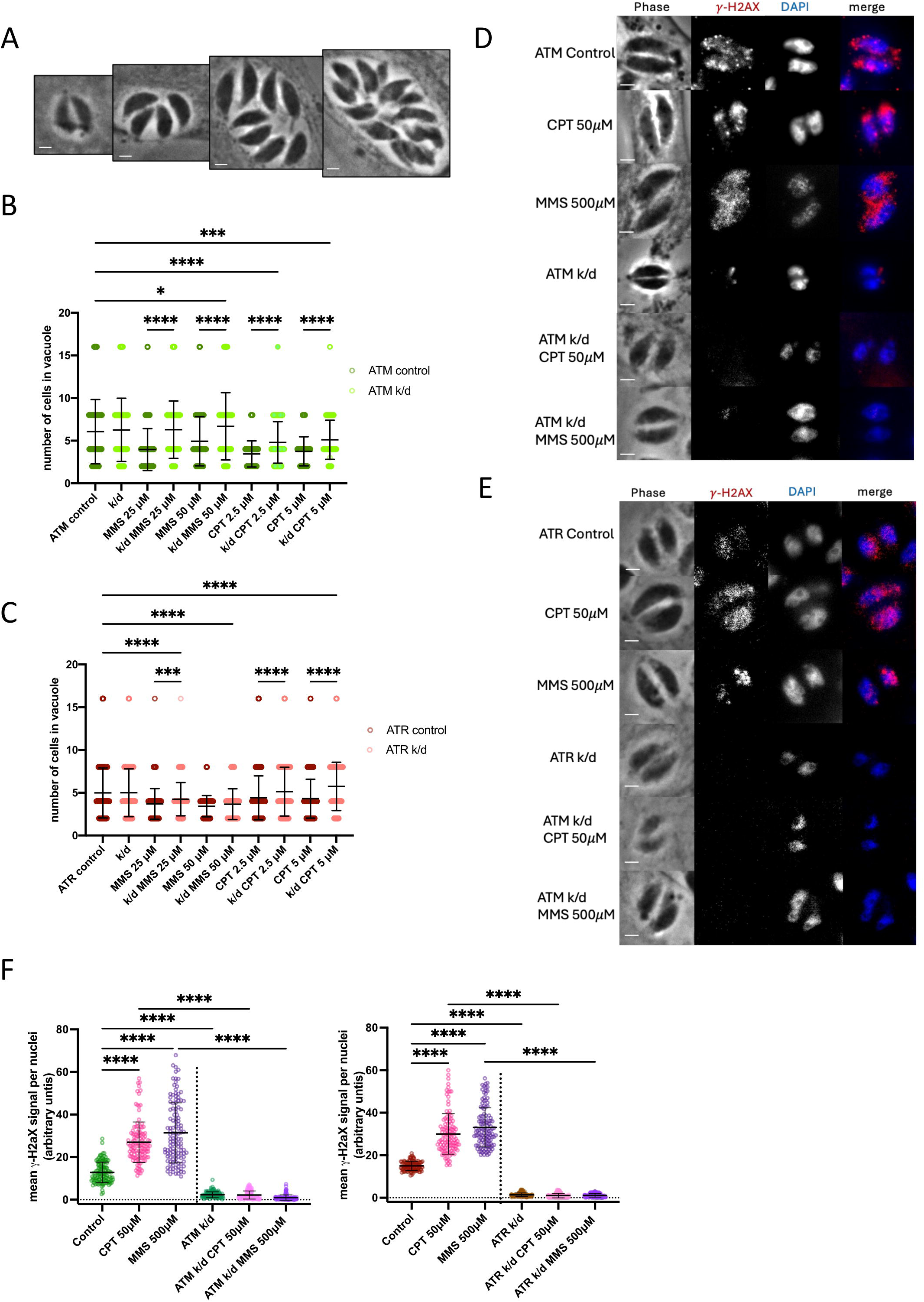
Knockdown of *Tg*ATM or *Tg*ATR ablates acute checkpoint-like responses to DNA damage. A. Images of representative vacuoles containing 2, 4, 8 or 16 parasites, as counted for the graphs in B and C after 24h growth. Scale bar 2 μm. B,C. Replication assays were conducted on *Tg*ATM k/d (B) and *Tg*ATR k/d (C) parasites, grown for 24h under DNA-damaging conditions. Numbers of parasites in 250 vacuoles were counted per condition in each experiment; data from 2-3 biological replicates are shown. Ordinary one-way ANOVA and post hoc Tukey’s multiple comparisons tests were made of control vs all other conditions, and of each drug treatment vs k/d with the same treatment. P values: *<0.05, **<0.01, ***<0.001, ****<0.0001. Comparisons not shown, ns. Also not shown on the graph, for concision: the ATM control line differed from the same line under all DNA damage conditions at p-value ****<0.0001; the ATR control also differed from all DNA damage conditions at p-value ****<0.0001 except ***<0.001 for 2.5uM CPT. D,E. IFA of H2AX phosphorylation in *Tg*ATM k/d (D) and *Tg*ATR k/d (E) tachyzoites that were treated for 2 h with DNA-damaging agents. Representative images from three biological replicates are shown. Scale bar 2 μm. F Mean H2AX signal in nuclei was quantified from representative images in D and E using ImageJ. 40 nuclei were quantified per condition for three biological replicates. Ordinary one-way ANOVA and post hoc Tukey’s multiple comparisons tests were performed for control vs k/d and single drug treatment; and single drug treatment vs k/d plus drug; P values: *<0.05, **<0.01, ***<0.001, ****<0.0001. Comparisons not shown, ns.

We then measured a second acute read-out for the DNA damage checkpoint: phosphorylation of histone H2AX. Foci of γH2AX appeared at substantial levels in normally replicating parasites, as previously reported [26–28], suggesting a high endogenous level of replication stress in cultured tachyzoites. Phosphorylation appeared to be induced even further by DNA damage, particularly via CPT, with brighter and larger foci of γH2AX appearing (Figure 4D, E). Both the individual knockdowns ablated the γH2AX signal almost entirely, and it could not be induced even after DNA damage (Figure 4F). Again, both proteins were apparently required, since either one of the individual knockdown affected H2AX phosphorylation equally severely.

### *Tg*ATR does not behave quantifiably differently in the presence or absence of *Tg*ATM

We wished to investigate the extent and nature of cooperation between *Tg*ATM and *Tg*ATR. Accordingly, we added a second, distinguishable tag (3x V5) to the ATR gene in the HA-tagged, inducible knockdown line for *Tg*ATM (Figure 5A). Thus, we could follow the location and quantity of *Tg*ATR in cells with *Tg*ATM either present or absent (the 250-650kDa size of these proteins precluding co-immunoprecipitation and western blotting to test for a direct interaction).

**Figure 5:**
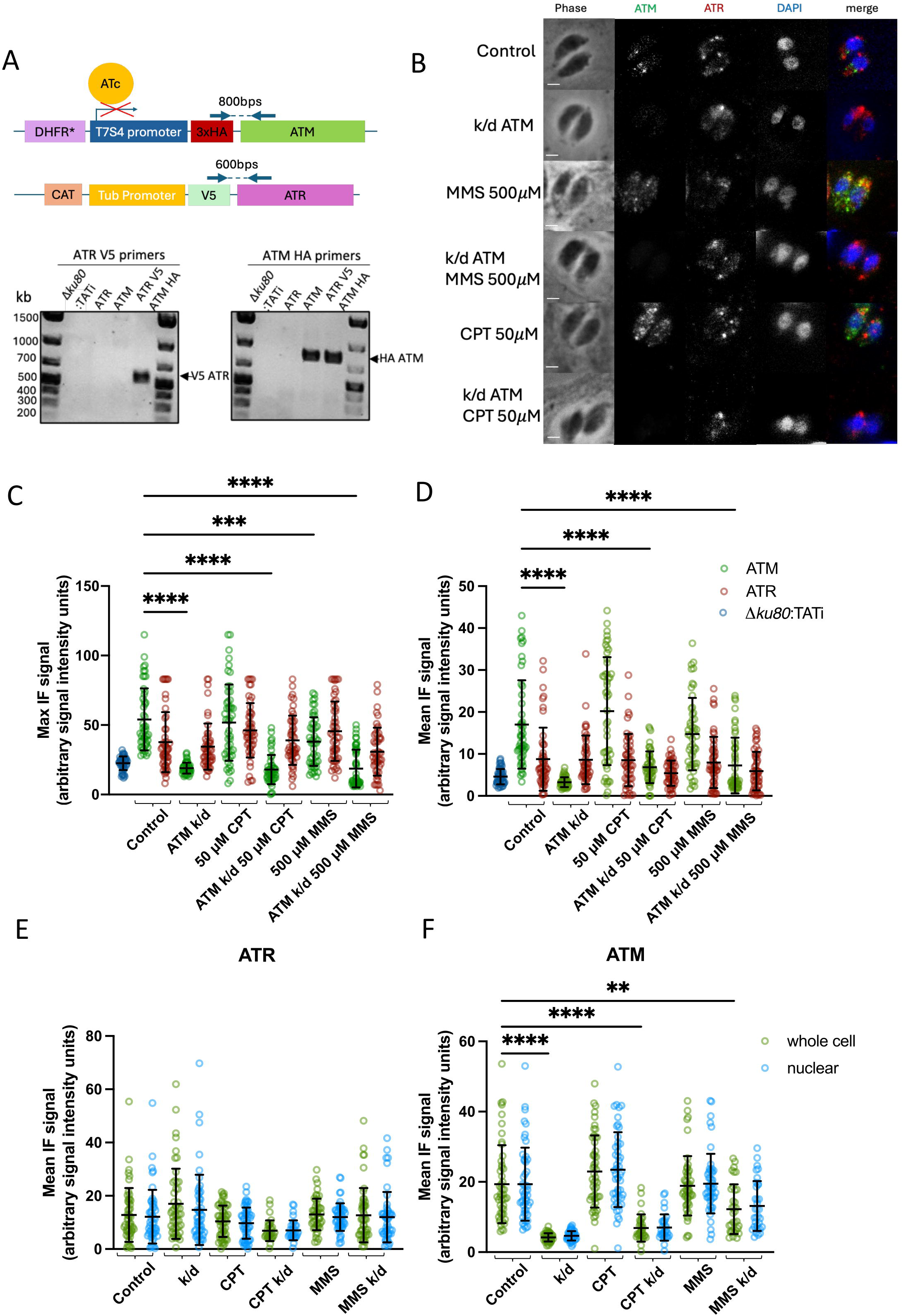
*Tg*ATR does not behave quantifiably differently in the presence or absence of *Tg*ATM. A. Construct used to tag the *TgATR* gene with V5, under the tubulin (Tub) promoter and chloramphenicol resistance selectable marker (CAT). Also shown is the original construct used to tag the *TgATM* gene with HA. Schematics show where PCR primers were located to test correct integration of the two tags. The PCR results on the left shows V5-tagged *TgATR* after transfection of this construct into the *Tg*ATM-HA line (no band appeared in the parental control line (‘Δ*ku80*:TATi’) or in either single-gene-tagged line); the PCR on the right shows that HA-tagged *TgATM* remained in the double-tagged line. B. IFA showing parasites with both ATR and ATM tagged, in either control or *Tg*ATM knockdown conditions, treated with DNA damaging drug treatments for 2 h, 24 h after infection of fresh host cells. Scale bar, 2 µm. C. Using IFA images as in (B), maximum brightness of *Tg*ATR and *Tg*ATM foci was measured in both control and ATM-knockdown conditions, as well as in the parental line (Δ*ku80*:TATi). One biological replicate was analysed; n = 50 parasites quantified per condition. Ordinary one-way ANOVA and post hoc Tukey’s multiple comparisons tests of control vs all conditions for both *Tg*ATR and *Tg*ATM were completed; P values: *<0.05, **<0.01, ***<0.001, ****<0.0001. Comparisons not shown, ns. D. Mean brightness of *Tg*ATR and *Tg*ATM signals throughout parasites, measured in both control and ATM-knockdown conditions, as well as the parental line (Δ*ku80*:TATi). N = 50 parasites quantified per condition. Statistical analysis as in (C). E,F. Using IFA images as in (B), the amount of signal in the nucleus and in the whole cell was quantified for both *Tg*ATR (E) and *Tg*ATM (F), with or without ATM k/d and addition of 50µM CPT or 500µM MMS. One biological replicate was analysed in this experiment; n = 50 parasites per condition. Ordinary one-way ANOVA and post hoc Tukey’s multiple comparisons tests of control vs all conditions for both *Tg*ATR and *Tg*ATM were completed; P values: *<0.05, **<0.01, ***<0.001, ****<0.0001. Comparisons not shown, ns.

Figure 5B shows that there was no substantial colocalisation of the two proteins: both appeared in speckles throughout the parasite that were largely non-overlapping. There was also no clear difference, detectable by immunofluorescence, in the location or amount of *Tg*ATR in the presence or absence of *Tg*ATM. The assessment of representative immunofluorescence images is inherently qualitative, so we also quantified the signals in cohorts of cells: both the amount of signal across individual parasites, and the maximum brightness of foci per parasite. There was considerable variation in both parameters, but ATR foci did not become reproducible fainter, brighter or more abundant in the absence of *Tg*ATM (Figure 5C, D). Therefore, although the two proteins were required in concert to induce checkpoint phenotypes (Figure 4), we could not detect any direct interaction between them. Finally, we checked whether the distribution of either protein changed after DNA damage, specifically to concentrate in the nucleus, as might be expected if a DNA damage response was required there. When the proportion of signal in the nucleus versus cytoplasm was quantified in cohorts of cells before or after DNA damage, no clear differences were seen (Figure 5E, F).

### A specific inhibitor of human ATM does not phenocopy the knockdown of *Tg*ATM or *Tg*ATR

Previous publications used the human ATM inhibitor KU-55933 on the assumption that it would be equally specific for *Tg*ATM [20, 21]. We tested this by comparing the effect of KU-55933 with the effect of knocking down *Tg*ATM. Figure 6A-C shows that the KU-55933 inhibitor caused a dose-dependent growth defect in a 10-day plaque assay, in contrast with either individual knockdown, which did not (Figure 3). Furthermore, the growth defect imposed by the inhibitor was similar whether an individual PIKK was also knocked down or not (Figure 6C). This led to two potential conclusions: either the effect of KU-55933 was ‘off target’ (via a pathway unrelated to *Tg*ATM/*Tg*ATR), or alternatively, the inhibitor was targeting both PIKKs and this complete loss of function was more deleterious for *T. gondii* than the loss of either one alone.

**Figure 6:**
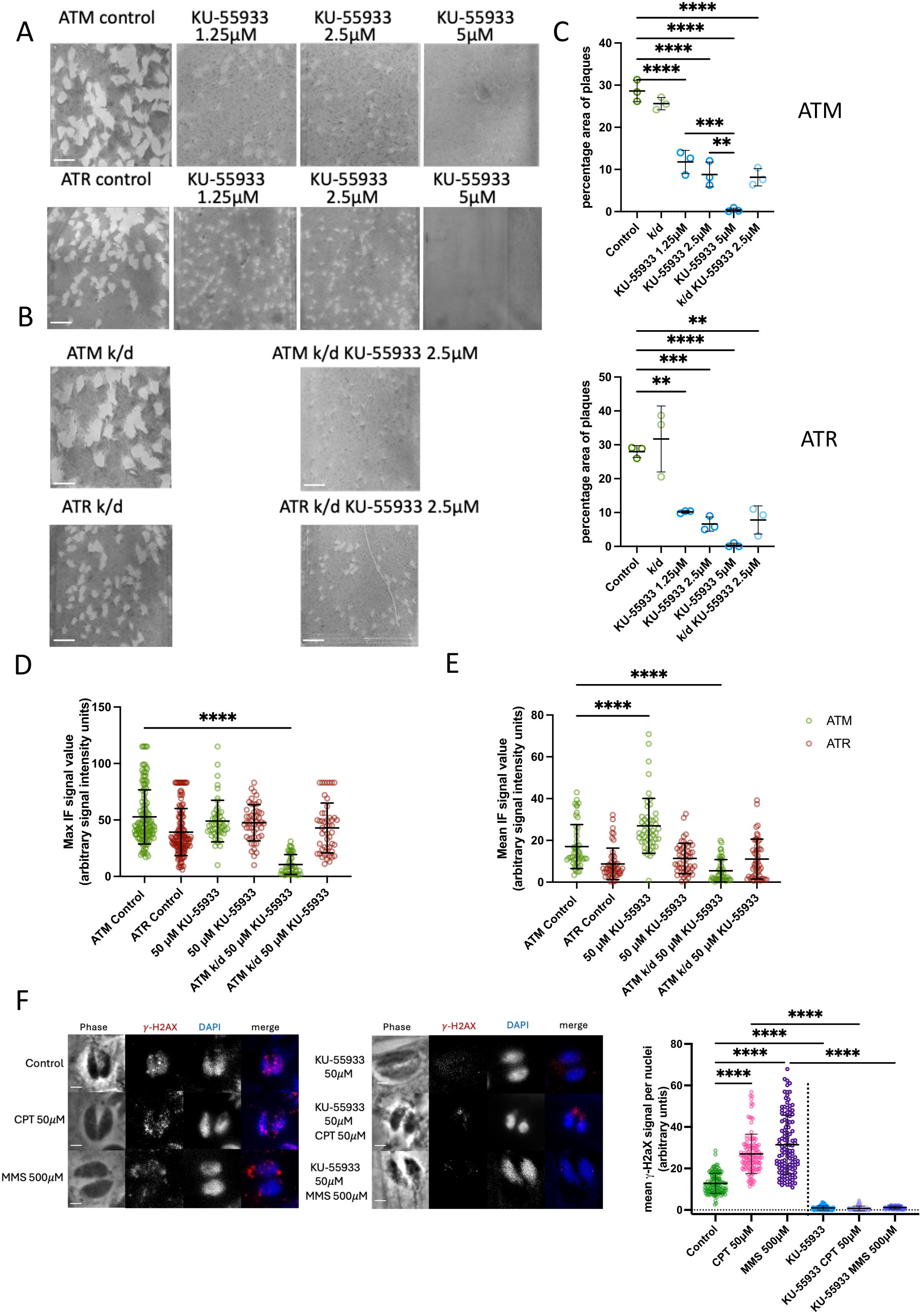
KU-55933, an inhibitor of human ATM, does not phenocopy knockdown of *Tg*ATM or *Tg*ATR. A. Images of plaque assays conducted in the *Tg*ATM and *Tg*ATR HA-tagged lines treated with increasing concentrations of KU-55933. B. Images of plaque assays showing the same lines as in (A), treated with KU-55933, after knockdown of *Tg*ATM or *Tg*ATR. C. The percentage area of plaques in each background of host cells was quantified from representative images shown in A and B: three biological replicates were quantified for each condition. Ordinary one-way ANOVA and post hoc Tukey’s multiple comparisons tests were made of control vs all, KU-55933 treatments against each other, and KU-55933 treatment at 2.5µM with vz without PIKK k/d. P values: *<0.05, **<0.01, ***<0.001, ****<0.0001. Comparisons not shown, ns. D. Using IFA images, the maximum brightness of *Tg*ATR and *Tg*ATM foci was measured in both control and ATM-knockdown conditions, with or without KU-55933. One biological replicate was analysed; n = 50 parasites quantified per condition. Ordinary one-way ANOVA and post hoc Tukey’s multiple comparisons tests of control vs all conditions for both *Tg*ATR and *Tg*ATM were completed; P values: *<0.05, **<0.01, ***<0.001, ****<0.0001. Comparisons not shown, ns. E. Mean brightness of *Tg*ATR and *Tg*ATM signals throughout parasites, measured in both control and ATM-knockdown conditions, with or without KU55933. N = 50 parasites quantified per condition. Statistical analysis as in (D). F. IFA showing H2AX foci in the parental strain, treated or not treated with KU55933, with or without DNA-damaging drugs MMS or CPT. Scale bar, 2 µm. Mean H2AX signals were then quantified in 40 nuclei per condition, from three biological replicates, and these are shown in the graph (control data are shown alongside, as in Figure 4). Ordinary one-way ANOVA and post hoc Tukey’s multiple comparisons tests were made; P values: *<0.05, **<0.01, ***<0.001, ****<0.0001. Comparisons not shown, ns.

We checked whether treatment with KU-55933 could affect either the amount of *Tg*ATM or *Tg*ATR in individual parasites, or the maximum brightness of their foci per parasite, by quantifying IFA images as in figure 5. No consistent changes were detected in the foci of either protein (except, as expected, when *Tg*ATM was knocked down) (Figure 6D). However, when the total quantity of protein per parasite was measured, KU-55933 treatment did correlate with an overall increase in the amount of *Tg*ATM, whereas there was no change in *Tg*ATR (Figure 6E). This implied that KU-55933 might indeed inhibit *Tg*ATM (and potentially also *Tg*ATR) and that the parasite might respond to this by upregulating *Tg*ATM expression.

To further distinguish between the possibilities of off-target killing or pan-PIKK inhibition, we checked the previously published result that cells treated with KU-55933 could not phosphorylate H2AX [21]. This was confirmed: the inhibitor largely prevented γH2AX foci from forming, both in endogenous replication and in response to DNA damage (Figure 6F). In this regard, inhibitor-treatment was similar to the individual knockdown of *Tg*ATM (or *Tg*ATR). Since KU-55933 inhibits human ATM as well, H2AX phosphorylation was equally inhibited in the human host cell nuclei, as expected (Figure S3). Overall, it appeared likely that KU-55933 did in fact inhibit *Tg*ATM, but that it additionally affected *Tg*ATR and/or other off-target proteins, resulting in a growth defect.

## DISCUSSION

This work characterised the two PIKK homologues in *Toxoplasma gondii*, with the aim of establishing whether or not they act as DNA damage checkpoint kinases. This was an important question because toxoplasmosis is often treated with drugs that may cause DNA damage, and also because it could help to establish how damage-responsive checkpoints have evolved across the apicomplexan phylum. In this large phylum of early-diverging protists, some parasites, like *T. gondii*, retain PIKK homologue(s) while other important parasites, like *Plasmodium* spp., appear to have lost them.

We established, phylogenetically, that *T. gondii* does encode orthologues of all the classes of PIKKs defined in human cells. We then tagged and knocked down the orthologues of ATM and ATR. The first striking observation was that neither protein was primarily nuclear, but appeared diffusely throughout the cytoplasm, in largely non-overlapping speckles that were inconsistent with any particular organelle. It is notable that another protein with a predicted nuclear role, the acetyltransferase *Tg*MYST, was also reported thus [20], and proposed to shuttle to the nucleus. In human cells, although ATM is nuclear, it can under some stimuli shuttle to the cytoplasm [29]. We, however, found no clear evidence that either *Tg*ATM or *Tg*ATR shuttled between cytoplasm and nucleus after DNA damage.

Both *Tg*ATM and *Tg*ATR were then inducibly knocked down, with near-100% efficiency as seen by immunofluorescence, and also by western blotting in the case of *Tg*ATM. As with any inducible gene regulation, some residual protein might remain. However, under severe-knockdown conditions, we found no evidence that either protein was essential or had any effect on normal cell growth. Nevertheless, after DNA damage, both knockdowns did ablate an acute reduction in cellular replication, as well as another key marker of the DNA damage response: phosphorylation of histone H2AX, which showed that a genuine kinase-driven checkpoint signal occurred, and that this required *Tg*ATM/*Tg*ATR. Thus, there was strong evidence for a canonical checkpoint role for these proteins. This was not initially detected in plaque assays simply because these assays occur over 10 days and many lytic cycles. Any parasites that failed to respond to DNA damage and continued to replicate would probably die from irreparable genome damage before a large plaque could be formed, resulting in a similar outcome to that seen in parasites that sustained damage and did make a checkpoint response, arresting or slowing their growth.

Our study did not address whether the DNA damage checkpoint in *T. gondii* is enforced equally in all phases of the cell cycle. In human cells, damage such as DSBs can be detected and repaired in G1 and G2 as well as S-phase, whereas other types of damage can be replication-dependent – detected efficiently only when replication forks are active during S-phase. Further dissection of this issue would require clear phase markers in *T. gondii:* for example, cells in S-phase might be detected by bromodeoxyuridine labelling of nascent DNA or fluorescent labelling of proliferating cell nuclear antigen (PCNA). These have both been achieved in *Plasmodium* [30, 31], but have yet to be demonstrated in *Toxoplasma*.

In human cells, ATR responds primarily to stalled replication forks and ATM, to DNA DSBs [32]. To establish whether *Tg*ATM and *Tg*ATR acted similarly, we used two different DNA damaging agents, the alkylating agent MMS, well-established to cause fork stalling at alkylated bases, and the topoisomerase inhibitor CPT, which can induce DNA breaks. (Of note, our knockdowns were made in a background deficient for non-homologous end-joining, but still competent to repair DSBs by homologous recombination.) Interestingly, we detected little ‘division of labour’ between *Tg*ATM and *Tg*ATR: the phenotypes in H2AX phosphorylation and cellular replication were similarly affected by both knockdowns. The only detectable difference was that high concentrations of MMS still elicited some replication-slowing when *Tg*ATR was lost, but not when *Tg*ATM was lost, and the converse was true of CPT, suggesting that *Tg*ATM may be more important in responding to alkylation damage and *Tg*ATR, to DNA breakage. This would be the opposite specialisation of protein function to that in human cells. Generally, however, both proteins were required to induce both types of DNA-damage checkpoint.

Despite this, we found no evidence for their colocalization or reciprocal up/down-regulation, so it is unlikely that the two proteins act in a stable complex. Mutual regulation, such as transphosphorylation, may be more likely. In human cells, ATM exists as an inactive dimer and is induced to autophosphorylate after DNA damage, whereupon it dissociates and its monomers act on downstream substrates [33]. Perhaps in *T. gondii*, activating phosphorylation event(s) occur primarily *in trans* and require both PIKKs, or perhaps both PIKKs act in mutually-essential ways on a single transducer downstream. (In humans, ATM and ATR activate Chk2 and Chk1 respectively [32], but neither of the Chk kinases has been identified in *T.gondii*). Biochemical experiments would be required to interrogate these hypotheses, such as generating complete phosphoproteomes of knockdown parasites, or co-immunoprecipitating the PIKKs to seek evidence for their cross-phosphorylation. With proteins of 250-650kDa, and of low abundance (neither was detected in the subcellular atlas of the *Toxoplasma* proteome [24]), such assays are non-trivial.

We attempted to further separate the activities of the two proteins using a ‘specific’ small-molecule inhibitor of human ATM, KU-55933, but this did not phenocopy the *Tg*ATM knockdown. The inhibitor severely affected cell growth, suggesting either off-target effects or an equal inhibition of both PIKKs, with this being more deleterious than either single knockdown. Since the inhibitor ablated H2AX phosphorylation, and had the same effect with or without either genetic knockdown, we favoured the latter explanation. Although we were unable to generate a double genetic knockdown, KU-55933 may itself generate this effect. If so, this implies that the complete loss of all PIKK activity does affect cell growth and viability. This would be expected, since DNA damage occurs during normal replication (as marked by γH2AX), and constant checkpoint activation is probably required in replicating tachyzoites. Curiously, *Tg*ATM and *Tg*ATR seem to be mutually required for this, but perhaps in each genetic knockdown, the single remaining protein achieves some level of self-activation (or a very low residual level of either knocked-down protein can transphosphorylate the other), thus yielding a low-level response and hence cell survival, whereas KU-55933-treated cells make no response at all.

Overall, we conclude that *T. gondii* does have PIKK kinases that enforce a DNA-damage checkpoint, slowing down cellular replication and phosphorylating the DNA damage marker H2AX. Unusually, both *Tg*ATM and *Tg*ATR seem to be required for this, regardless of the type of DNA damage. This could represent a step towards the tendency for reductive evolution of PIKKs in apicomplexans: ATR is found broadly, although lost from *Plasmodium* spp., whereas ATM is independently lost from *Cryptosporidium*, *Babesia* and *Theileria* (Figure 1). This might, in turn, help to explain the apparently-unique situation in *Plasmodium*, where both PIKK homologues are entirely lost. (Whether checkpoint activity is then lost in *Plasmodium*, or transferred to another protein, remains to be explored.)

In terms of cell division, apicomplexan parasites are particularly interesting because they can divide by different modes at different lifecycle stages. For example, *Plasmodium* uses schizogony in the blood stage and something resembling endopolygeny in the oocyst [34]; *T. gondii* uses endodyogeny in intermediate hosts and a division mode that is variously defined in the literature as endopolygeny or schizogony in the definitive feline host [35–38]. Whether DNA damage checkpoints are equally active in both stages cannot presently be investigated, because experimental access to the feline stage in *T. gondii* is very limited. Nevertheless, further investigation of cell-cycle checkpoints in *T. gondii* might usefully inform the study of checkpoints in other apicomplexans such as *Plasmodium* as well.

## METHODS

### Database sampling and phylogenetic analysis

A custom database was collated that included data for relevant eukaryote species from UniProt [39] and the Marine Microbial Eukaryote Transcriptome Sequencing Project [40], as well as data for apicomplexans and close relatives from VEuPathDB [41] and two previous studies [42, 43]. This custom database was then sampled for ATM, ATR and TOR homologues using the program blastp (BLAST+ version 2.11.0 [44]), for which characterised homologues of these proteins from selected model organisms, including *T. gondii*, were used as search queries. The dataset of possible ATM, ATR and TOR homologues obtained was then clustered with CD-HIT [45] to remove highly similar sequences, thereby reducing overall redundancy of the dataset. Iterative rounds of alignment were then performed using mafft (MAFFT version 7.475 [46]), conserved site selection using trimal with “gappyout” mode (trimAl version 1.4 [47]), and tree inference with FastTreeMP (FastTree version 2.1.11 [48]) using the default settings and manual removal of sequences to further reduce dataset redundancy. Manual removal also identified and removed sequences that aligned poorly or were so dissimilar from the majority of the sequences in the dataset that they were unlikely to be ATM, ATR or TOR homologues, but rather false positives of the described sampling strategy. The final curated dataset that these methods obtained was then aligned using mafft-linsi (MAFFT version 7.475 [46]) and conserved sites were selected using trimal with “gappyout” mode (trimAl version 1.4 [47]). A phylogeny was then inferred from this dataset using the program iqtree2 (IQ-TREE version 2.1.2 [49]) with 1000 ultrafast bootstrap replicates (UFBoot2 [50]) and using the best-fitting model, LG+F+I+G4 ([51]) that was chosen according to the Bayesian Information Criterion by ModelFinder [52], all implemented within iqtree2. The protein sequences, alignment, and tree inference output files used to generate the phylogeny are provided at [Journal preferred repository?].

### Parasite strains and growth

*Toxoplasma gondii* (*T. gondii*) tachyzoites from the strain RH and derived strains, including Δ*ku80*:TATi tachyzoites (a gift from Lilach Sheiner and Boris Striepen, University of Georgia) [53], were grown on a confluent monolayer of human foreskin fibroblast (HFF) cells in 10 ml flasks of ED1 (Dulbecco’s Modified Eagles Medium supplemented with 1% foetal bovine serum (FBS), 0.2 mM L-glutamine, 0.25 μg ml^−1^ Fungizone (ThermoFisher), 10 unit ml^−1^ penicillin and 10 μg ml^−1^ streptomycin (ThermoFisher)) and maintained at 37 °C with 10% CO_2_, as previously described [54].

### Creation of transgenic *T. gondii*

Gene tagging was performed as described by Waller and colleagues [23]. A CRISPR/Cas9-assisted, PCR-mediated tagging strategy was used to perform endogenous gene tagging with epitope tags and/or endogenous promoter replacement with an ATc-regulatable t7s4 promoter. The generation of transgenic *T. gondii* involved a level M Cas9/sgRNA plasmid, P5, containing a specific protospacer sequence for directing a locus-specific DNA break to facilitate homologous recombination-driven insertion of the donor DNA fragment. Protospacer sequences were designed for insertion into the N-terminus of genes encoding putative *T. gondii* homologues of ATR (TGGT1_283702) and ATM(TGGT1_248530) (Figure S4). The donor DNA fragment was PCR amplified from one of the template plasmid vectors (Figure S4A) using primers (Figure S4B) that included specific homology arms directing the integration of this donor DNA construct into the target genetic locus for ATM and ATR. Electroporation of these constructs was completed as in [23].

### Parasite genomic DNA extraction for PCR confirmation

T25 flasks containing mostly extracellular *T. gondii* were scraped and host cell debris was pelleted at 100 *xg* for 5 min. A 1-1.5 ml aliquot was harvested from the supernatant, containing 0.5-5 x 10^6^ fully lysed *T. gondii* tachyzoites, and these were collected by centrifugation (7,000 *xg*/full speed, 2 min). The cell pellet was resuspended in 20 μl proteinase K solution: 5% proteinase K (Thermo Scientific, EO0491), 10% PCR buffer 10x (Roche, 11732650001), 85% dH_2_O. This was incubated at 45 °C for 60 min. The proteinase was heat inactivated at 94 °C for 10 min. The remaining solution was used to run PCR directly, 0.3 - 1.5 µl as DNA template for a 20 µl PCR reaction.

### Plaque assay

Plaque assays required parasites derived from freshly lysed host cells, which were centrifuged at 100 *xg* for 5 min to remove host cell debris. The same number of parasites (around 500) was added to each new T25 flask containing HFF monolayer. In experiments testing the knockdown (k/d) of a target gene, a control of untreated *T. gondii* infected HFFs was used, as well as ATc-treated (k/d) *T. gondii* infected HFFs. For the ATc treatment, 0.5 μg/ml of ATc was added. After 10 days of incubation at 37 °C and with 10% CO_2_, the media from flasks was aspirated and the cells were washed once with PBS. Cells were fixed with 5 ml of 100% ice-cold methanol for 5 min and stained with 5 ml of 1% crystal violet solution for 15 min. After staining, crystal violet solution was removed and flasks were washed three times with PBS. Flasks were allowed to dry, then imaged for qualitative analysis. Quantitative analysis was also performed on representative fields from three biological replicates, calculating the percentage plaque area in each field of host cells via ImageJ.

### Immunofluorescence

*T.gondii* infected HFFs were grown on glass coverslips in 6 well plates. Cells were fixed to coverslips with 2% PFA/PBS for 15 min at RT and permeabilised with 0.02% TritonX100 in PBS for 10 min. Cells were blocked for 1 h in 2% BSA/PBS. Coverslips were incubated with primary antibody in blocking solution for 1 h and then incubated in secondary antibodies in 2% BSA/PBS for 1 h in the dark. 120 μl of 2 μg/ml DAPI solution (Merck) was added for 10 min and then coverslips were mounted on slides in ProLong® Diamond Antifade (ThermoFisher, P36961). Images were acquired using a Nikon Eclipse Ti widefield microscope with a Nikon objective lens (Plan APO, 100x/1.45 oil), and a Hamamatsu C11440, ORCA Flash 4.0 camera. Images were processed using the NIS-Elements software and ImageJ (v.1.51w).

### Replication assay

For the replication assay parasites were pre-treated with or without ATc for 48 h. Parasites derived from freshly lysed host cells were counted and an equal number (1.6 x 10^6^) was added to HFF monolayers grown on coverslips in 6-well plates. Cells were incubated for 2 h at 37°C to allow invasion. Parasites that had not invaded were washed off three times with warm ED1. Parasites were then exposed to the DNA damaging agents CPT or MMS, allowed to grow for a further 24 h (still with or without ATc), then fixed as per the protocol for immunofluorescence and mounted onto slides using DAPI stain only. Images were acquired using a Nikon Eclipse Ti widefield microscope with a Nikon objective lens (Plan APO, 100x/1.45 oil), and a Hamamatsu C11440, ORCA Flash 4.0 camera. Images were processed using the NIS-Elements software and ImageJ (v.1.51w). Numbers of parasites in each vacuole were counted for 250 vacuoles per condition.

### Immunoblotting

Approximately 50 × 10^6^ parasites were purified from the host cell debris by filtration through 3-μm-pore-size polycarbonate film membrane filters (Nuclepore Track-Etch Membrane, Whatman). Cells were collected and washed in PBS by centrifugation at 1,700 × *g* for 10 min at RT, then pellets were resuspended in 1x RIPA buffer (150 mM sodium chloride, 1.0% NP-40 or Triton X-100, 0.5% sodium deoxycholate, 0.1% SDS (sodium dodecyl sulfate) and 50 mM Tris pH 8, added to 50 ml dH_2_0). Protease inhibitors (ROCHE, cOmplete Mini) were added to 10 ml aliquots directly before use). Samples were freeze-thawed 3 times by placing tubes in dry ice / 100% ethanol and a heated Eppendorf block at 37 °C. Cell debris was pelleted at 4 °C at 13,000 *xg*, then supernatant was removed to a new Eppendorf. 1 x NuPage LDS Sample Buffer (ThermoFisher: NP0007), supplemented with 0.1% 2-mercaptoethanol, was added to samples and boiled at 90 °C for 5 min. Protein lysates were resolved on SDS-PAGE in a 4-15% TRIS-glycine gel (BIO-RAD Mini-PROTEAN® TGX™) together with a molecular weight marker, Page Ruler Plus (ProteinTech). Gels were run for 1–2 h at 100 V until the ladder 10 kDa band was at the bottom of the gel. Nitrocellulose membrane 0.2-μM-pore-size (Amersham™ Protran®, GE Healthcare) was cut to size and wetted in transfer buffer. Wet electro-transfer was completed in transfer buffer at 60 V for 45 min.

Membrane was blocked with 3% BSA TBST for 1 h, washed once in TBST and incubated overnight in primary antibody diluted in blocking buffer at 4 °C. The blot was washed 3x 5 min in TBST and incubated in a secondary antibody diluted 1:5000 in blocking buffer for 1 h. Blots were washed again 3x 5 min, TBST, developed using SuperSignal West Pico Plus Chemiluminescent Substrate (Thermo Scientific) and imaged on the gel imager (Azure, Q500).

### IFA quantification analysis

IFA images were analysed by creating regions of interest (ROI) encompassing single parasites, or nuclei, in Image J. ROIs were created in ImageJ using the ‘threshold, analyse particles’ function. ROIs of between 500-5000 pixels were selected; overlapping cells and top-down cells were removed visually. For nuclei vs whole cell analyses, ROIs for the nuclei were made using DAPI channel images and then matched to whole cell ROIs using X, Y coordinates. Mean (average signal per cell), and Max (maximum signal pixel per cell) gray values were measured from each channel. One biological replicate was analysed for these analyses.

## ACKNOWLEDGEMENTS

This work was funded by the European Research Council (ERC) under the European Union’s Horizon 2020 research and innovation programme (ERC-2016-COG 725126 to CJM); Wellcome Award 214298/Z/18/Z to RFW; and a PhD studentship from the University of Cambridge Department of Pathology to MJK. The funders had no role in study design, data collection, interpretation or the decision to submit the work for publication.

## SUPPLEMENTARY FIGURE LEGENDS

Figure S1: <u>Homologues of cell cycle checkpoint kinases in Apicomplexans and other eukaryotes</u>

A fully expanded version of the phylogenetic tree shown in Figure 1.

Figure S2: <u>Inducible knockdown of *Tg*ATM and *Tg*ATR</u>

*A.* The phenotype score for *Tg*ATM from a genome-wide CRISPR screen to identify essential genes in *T. gondii* (Sidik et al., 2016).

*B.* The phenotype score for *Tg*ATR from a genome-wide CRISPR screen to identify essential genes in *T. gondii* (Sidik et al., 2016).

*C.* The schematic shows where the primers were located for PCR-based testing of correct integration of the HA tag on the *ATM* gene. The PCR shows the correct size band was present in all ATM clones and not in the parental control line. Asterisks denote non-specific bands that amplified from both the parental line (‘Δ*ku80*:TATi’) and HA-tagged clones.

*D.* The schematic shows primers designed to test for correct integration of the HA tag on the *ATR* gene. The PCR shows correct integration in several clones and the absence of a product from the parental control line; an HA-tagged ATM control (as in C) is also shown.

*E.* Anti-HA western blot of samples from an ATM tagged clone, taken at 24, 48 and 72 h after the addition of ATc. Profilin was used as a control protein.

Figure S3: <u>ATc has no effect on the DNA damage checkpoint response of the host cell of</u> *<u>T. gondii</u>*<u>, whereas KU-55933 inhibits this response.</u>

Immunofluorescence images showing that phosphorylation of H2AX in the host cell nucleus a) was not affected by addition of ATc, in cells either uninfected or infected with the parental line of *T. gondii*, Δ*ku80*:TATi; b) was inhibited upon addition of the KU-55933 inhibitor. Cells were treated with 0.5mg/ml ATc, 50 μM CPT, 50μM KU-55933, or combinations of these. Representative images from one biological replicate are shown. Scale bar 5 μm.

Figure S4: <u>Resource table</u>

*A.* Table containing plasmid maps.

*B.* Table containing protospacer and primer sequences.

*C.* Table containing antibodies used.

