## Supplementary figures and images for "Characterisation of cell cycle checkpoint kinases in Toxoplasma gondii"

### Fig S1

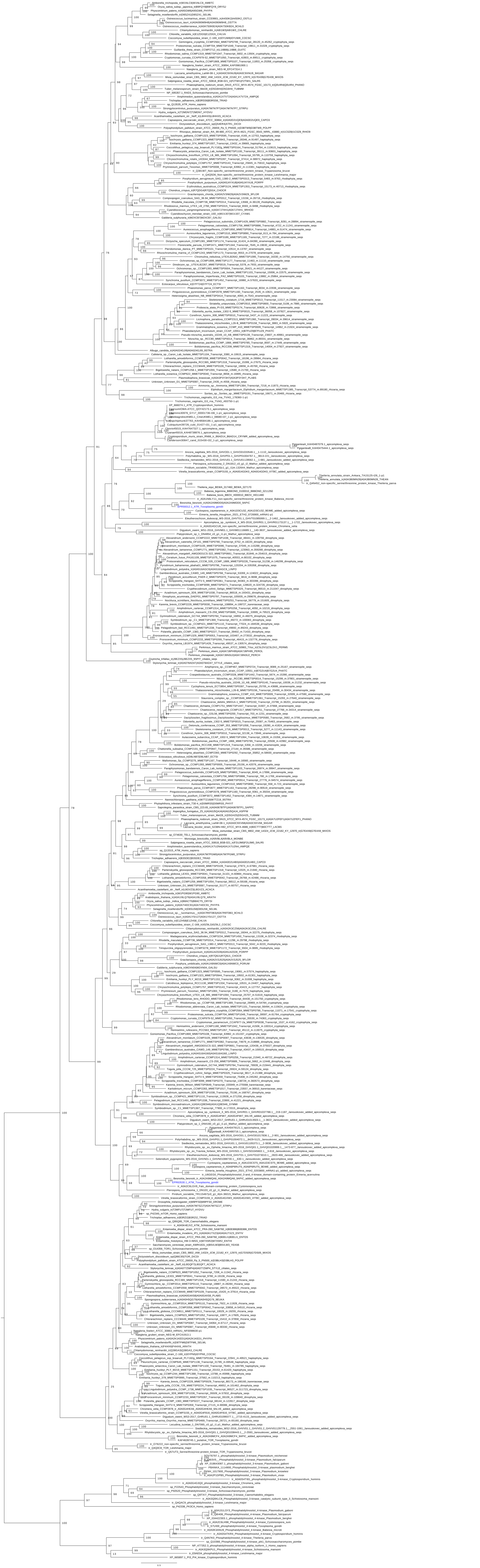
