## Supplementary material for "Characterisation of cell cycle checkpoint kinases in Toxoplasma gondii": Fig S2-4

Figure S2:

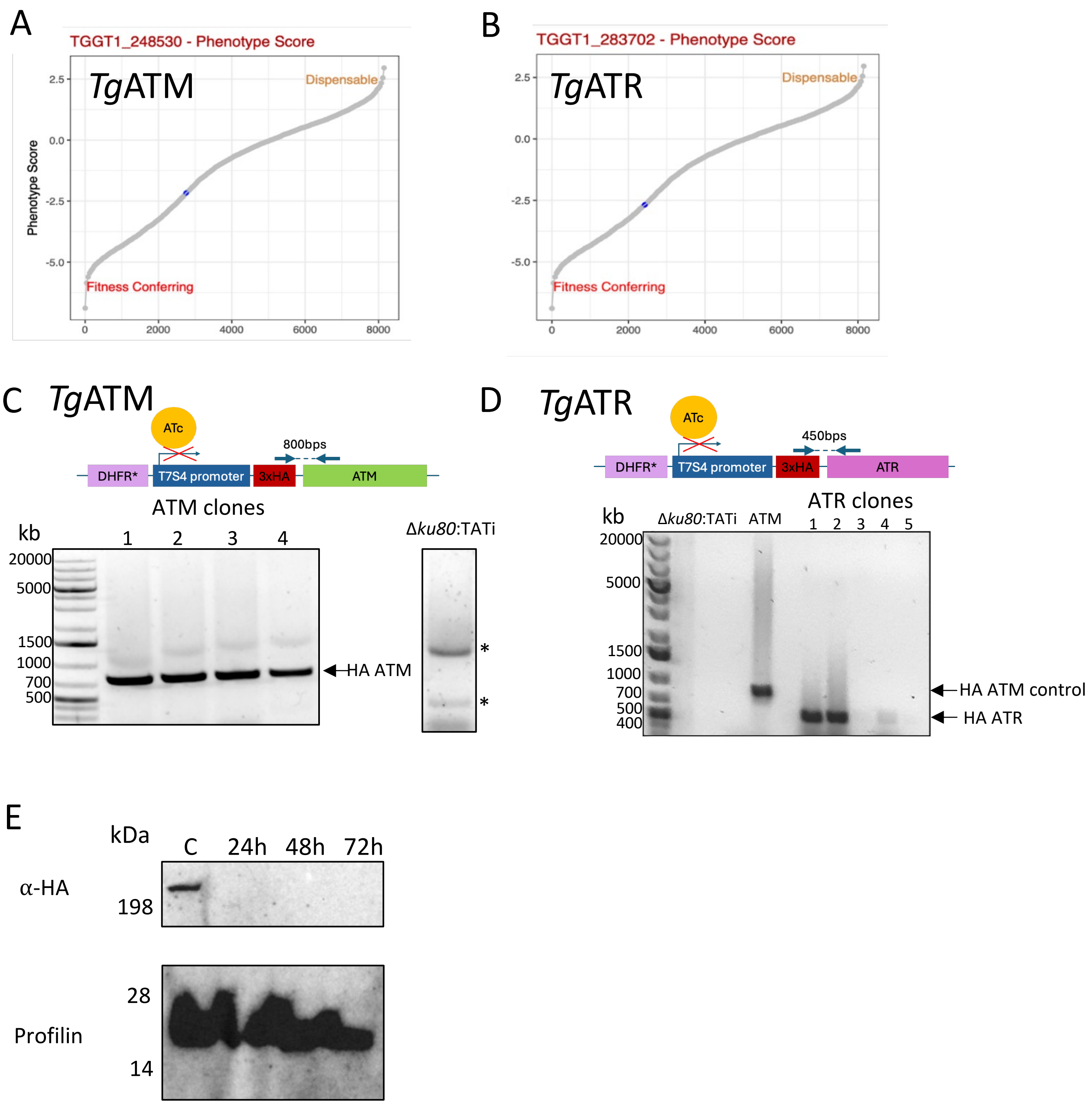

A
TGGT1_248530 - Phenotype Score
TgATM
Dispensable
Fitness Conferring
Phenotype Score
2.5
0.0
-2.5
-5.0
0
2000
4000
6000
8000
B
TGGT1_283702 - Phenotype Score
TgATR
Dispensable
Fitness Conferring
C
TgATM
ATc
800bps
DHFR*
T7S4 promoter
3xHA
ATM
ATM clones
kb
1
2
3
4
Δku80:TATi
20000
5000
1500
1000
700
500
HA ATM
*
*
D
TgATR
ATc
450bps
DHFR*
T7S4 promoter
3xHA
ATR
kb
Δku80:TATi
ATM
ATR clones
1
2
3
4
5
20000
5000
1500
1000
700
500
400
HA ATM control
HA ATR
E
kDa
C
24h
48h
72h
α-HA
198
28
Profilin
14

Figure S3:

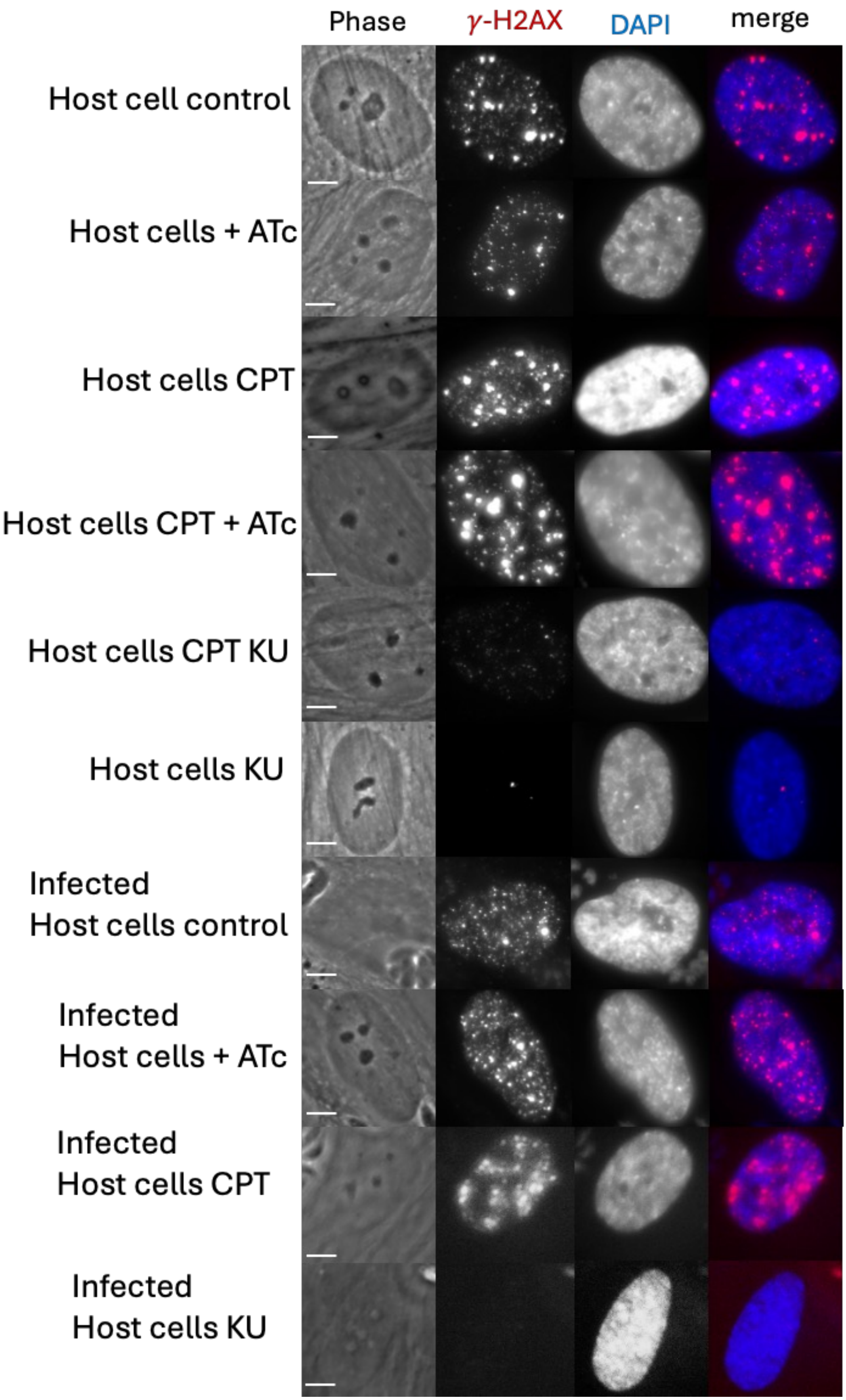
Phase
γ-H2AX
DAPI
merge
Host cell control
Host cells + ATc
Host cells CPT
Host cells CPT + ATc
Host cells CPT KU
Host cells KU
Infected
Host cells control
Infected
Host cells + ATc
Infected
Host cells CPT
Infected
Host cells KU

### Figure S4A:

<table>
<tr><th>Resource</th><th>Use</th><th>Sequence</th><th>Source</th></tr>
<tr><td>Level M CRISPR/Cas9 P5</td><td>To transfect T. gondii and make transgenic lines, by cutting ATR and ATM and sgRNA sites.</td><td>Golden Gate Level M plasmid containing genes for expression of the protospacer-sgRNA (PS-sgRNA) sequence under a T. gondii U6 promoter, the Cas9-HA- GFP fusion protein under a T. gondii Sag1 promoter. Confers an ampicillin resistance cassette for the bacterial selection.<br>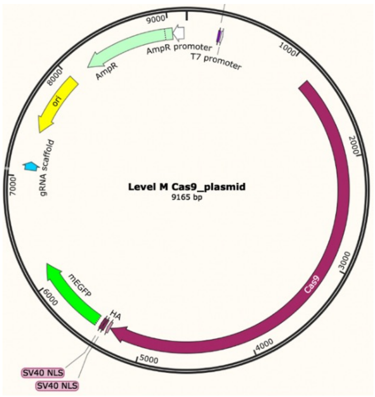
</td><td>(Barylyuk et al., 2020)</td></tr>
<tr><td>pPR2-3HA plasmid</td><td>Template for N-terminal 3x HA tagging and replacing the endogenous promoter to the t7s4 promoter</td><td>DHFR promoter-DHFR-DHFR terminator with t7s4 promoter-3x HA. Confers resistance to pyrimethamine in T. gondii.<br>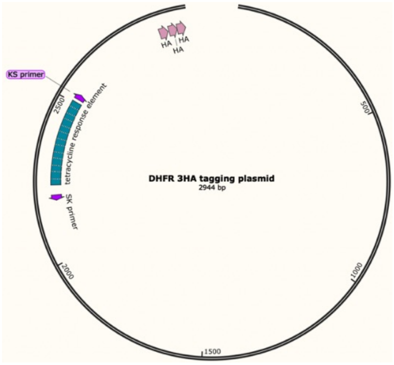
</td><td>(Katris et al., 2014)</td></tr>
<tr><td>pNter-CAT-3V5</td><td>Template for N-terminal 3x V5 tagging</td><td>Gra promoter-CAT-Sag1 with Tub promoter-3xV5-linker<br>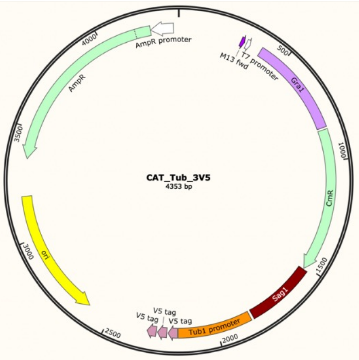
</td><td>Generated by Dr Ludek Koreny</td></tr>
<tr><td>DHFR terminator plasmid</td><td>Plasmid template for amplifying</td><td>Gra-CAT-Sag1; Tub1 Promoter-eGFP-linker-K13-DHFR terminator; UPRT 3' flank; endlinker pos3 in Level M<br>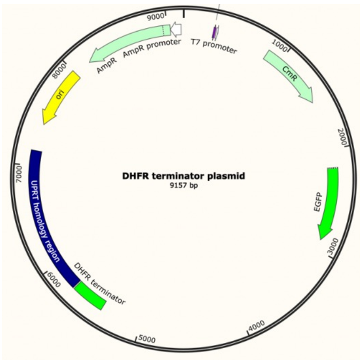
</td><td>Generated by Dr Ludek Koreny</td></tr>
</table>

Figure S4B:

| **Resource** | **Use** | **Sequence** | **Source** |
|---|---|---|---|
| Protospacer sequences | | | |
| ATR protospacer PS-sgRNA | ATR Protospacer | ATTGAACCTGCCGAGACAGT NGG | Designed by Sara Chelaghma |
| ATM protospacer PS-sgRNA | ATM protospacer | GACGACGAAAGAATGATCTT NGG | Designed by Dr Ludek Koreny |
| Primers | | | |
| 283702_sgRNA_5endF | To incorporate ATR PS-sgRNA | TGTGGTCTCAAGTTGATTGAACCTGCCGAGACAGTGTTTTAGAGCTAGAAATAGCAAG | Designed by Sara Chelaghma |
| ATM_sgRNA_5end_F | To incorporate ATM PS-sgRNA | TGTGGTCTCAAGTTGACGACGAAAGAATGATCTTGTTTTAGAGCTAGAAATAGCAAG | Designed by Dr Ludek Koreny |
| Universal_sgRNA_Rv | Amplify protospacer-sgRNA | TGTGGTCTCAAGCGGAAAAAGAAAAAAAAAGCACCGACTC | (Barylyuk *et al.*, 2020) |
| 283702_T7S4_F | ATR 3HA ts74 tagging | GTGCATGCTTTGTGCCGCTGCTCCACCAACTCGTTGTCGAGACAACTCTGTC | Designed by Sara Chelaghma |
| 283702_T7S4_R | | GCATGCCCCGCAAAGCAAACCAAACAGAGCGTCATTCCAGATCCTCCGGCATAATCTGGAACATCGTAAGGA | Designed by Sara Chelaghma |
| ATM T7S4 rv | ATM 3HA ts74 tagging | GGCAGGTGATACGAATGAAGGTGAACCGAAGATTCCAGATCCTCCGGCATAATCTGGAACATCGTAAGG | Designed by Dr Ludek Koreny |
| ATM T7S4 fw | | CCGGGAACCGGAATGAGCGGAGAAACAAAGACGACGAAAGACGTTGTCGAGACAACTCTGT | |
| 283702_N_F | V5 tagging of ATR | GTGCATGCTTTGTGCCGCTGCTCCACCAACTCAGGTCTCATGCCGGAG | Designed by Sara Chelaghma |
| 283702_N_R | | GCATGCCCCGCAAAGCAAACCAAACAGAGCGTCATAAATTCTCCAGATCCTGCAG | |
| ATR confirming PCR rv | Primers to confirm ATM and ATR 5' tags | CCTTCGACTGTCTCAC | Designed by Monique Johnson |
| ATM confirming PCR rv | | AGGGATACAGAAATCTGC | Designed by Monique Johnson |
| HA primer fw | | TACCCGTACGACGTC | Designed by Monique Johnson |
| V5 tag primer fw | | CCTATTCCCAATCCTCTTC | Designed by Monique Johnson |

Figure S4C:

| Antibodies | |
|---|---|
| Primary antibodies | |
| mouse α-V5 | (Invitrogen, R960-25) |
| rat α-HA | (ROCHE, 11867423001), |
| rabbit α-γ-H2aX | (Cell signalling, 9718S) |
| Secondary antibodies | |
| AlexaFLUOR™ 594 goat α-mouse IgG2a | (Invitrogen, 2044860) |
| AlexaFLUOR™ 488 Goat α-rat IgG (H+L) Cross-Adsorbed Secondary Antibody | (Invitrogen, 2551392) |
| AlexaFLUOR™ 647 goat α-mouse and goat α-Rabbit IgG (H+L) Cross-Adsorbed Secondary Antibody | (Invitrogen 21235) |
| AlexaFLUOR™ 647 goat α-Rabbit IgG (H+L) Cross-Adsorbed Secondary Antibody | (Invitrogen 21244) |
